# Cross-talk of cellulose and mannan perception pathways leads to inhibition of cellulase production in several filamentous fungi

**DOI:** 10.1101/520130

**Authors:** Lara Hassan, Liangcai Lin, Hagit Sorek, Thomas Goudoulas, Natalie Germann, Chaoguang Tian, J. Philipp Benz

## Abstract

It is essential for microbes to acquire information about their environment. Fungi use soluble degradation products of plant cell wall components to understand the substrate composition they grow on. Individual signaling pathways have been well described. However, the interconnections between pathways remain poorly understood. In the present work, we provide evidence of “confusion” due to cross-talk between the perception pathways for cellulose and the hemicellulose mannan in several filamentous fungi, leading to the inhibition of cellulase expression. We used the functional genomics tools available for *Neurospora crassa* to investigate this signaling overlap at the molecular level. Cross-talk and competitive inhibition could be identified both during uptake by cellodextrin transporters and intracellularly. Importantly, the overlap is independent of CRE-1-mediated catabolite repression. These results provide novel insights into the regulatory networks of lignocellulolytic fungi and will contribute to the rational optimization of fungal enzyme production for efficient plant biomass depolymerization and utilization.

## Introduction

Fungi are of ecological, economical, pharmaceutical and biotechnological importance. This group of microorganisms has a major commercial impact in product areas including food and feed, pulp and paper, textiles, detergents, bio-fuel and chemical production [1]. The importance of filamentous fungi in biotech lies in their potential to efficiently degrade plant cell wall material and release sugar monomers [2]. They utilize their cellular resources for the production of a wide range of enzymes including cellulases and hemicellulases. The great heterogeneity and resulting chemical complexity of lignocellulosic feedstocks provides a range of fermentable carbohydrates for high value biological, chemical, and pharmaceutical products [3]. Yet, the production of the fungal cellulolytic and hemicellulolytic enzymes for hydrolysis of complex biomass remain a high cost factor [4]. Research efforts to optimize enzyme production and remove unwanted constraints therefore are still warranted. Previous research has greatly focused on how filamentous fungi regulate the degradation of single polysaccharides as isolated cell wall components. However, relatively little is known about the cross-talk between separate signaling pathways for cellulose and hemicellulose perception during the utilization of complex carbon sources. In this study, we demonstrate that cross-talk not only occurs but can result in inhibition with detrimental effects for the production of hydrolytic enzymes.

Cellulose and hemicellulose are the major constituents of lignocellulosic biomass. While cellulose is a linear chain of glucose molecules connected by β-(1,4)-glycosidic linkages [5], hemicelluloses are a heterogeneous group of branched and linear polysaccharides [6] consisting mainly of xylans and mannans in variable ratios depending on the source of the biomass. While xylans, such as glucuronoxylan, arabinoxylan, and arabinoglucuronoxylan [7], are the most abundant hemicellulose in hardwoods, glucomannan represents the major hemicellulose in softwood (15%-20%) [8]. It consists of a β-(1,4) linked D-mannopyranose and D-glucopyranose backbone in a Man:Glc ratio of about 1.6:1 [9]. Cellulose and glucomannan are hydrolyzed by glucanases and mannanases into cello- and (gluco-)mannodextrins, respectively, which are further processed into the simple constituent monosaccharides by intra- and extracellular β-glucosidases and β-mannosidases [10,11]. The production of such enzymes is controlled by complex signaling networks including several transcriptional regulators. In *N. crassa*, CLR-1 and CLR-2 (cellulose degradation regulator 1 and 2) are essential transcription factors (TFs) responsible for the vast majority of the cellulolytic response [12]. In the presence of cellulose or its degradation products (such as cellobiose) as an inducer [10], a signaling pathway results in the activation of CLR-1 which in turn induces the expression of β-glucosidases and the cellodextrin transporter-encoding genes *cdt-1* and *cdt-2*. Both CDT-1 and CDT-2 are Major Facilitator Superfamily (MFS)-type transporters described to be capable of transporting cellobiose/cellodextrins into the cell [13]. Additionally, CLR-1 induces the expression of the transcription factor CLR-2, which in turn triggers the major cellulolytic response [14].

Homologs of these regulators are present in most filamentous Ascomycetes, albeit differing in their functional role [15–17]. For example, ManR, the CLR-2 ortholog in *Aspergillus oryzae*, is involved in the regulation of both cellulolytic and mannanolytic genes [18], a function that is partly conserved in *N. crassa* [14,19], while the function of the CLR-2 homolog in *Trichoderma reesei* (TR_26163) for the production of cellulase and hemicellulase is less clear so far [20]. In *T. reesei* and *Aspergillus* spp., the regulator XYR1/XlnR controls both the hemicellulolytic and the cellulolytic response [21–24] which is divergent from the mechanism utilized by *N. crassa*. The XYR1-homolog in *N. crassa*, XLR-1, is more specific for the regulation of hemicellulose degradation, yet it only modulates cellulase induction [25]. In the presence of a preferred carbon source, another highly conserved regulatory system, carbon catabolite repression (CCR), is activated to repress unnecessary metabolic routes and prevent the wasting of energy. A key component of CCR in filamentous fungi is the TF CreA/CRE1/CRE-1, which represses the expression of genes encoding enzymes involved in lignocellulose degradation [26–30]. The presence of partially conserved regulatory mechanisms for lignocellulose degradation [17,31] and the partially different functions assigned to homologous regulators in the various fungal species add another level of complexity to the regulation of lignocellulolytic genes. However, the elucidation of the underlying mechanisms in those fungi, despite (or precisely because of) existing differences and similarities, is likely the key to a better understanding of how fungi utilize transcriptional rewiring to enable efficient plant biomass degradation adapted to their specific ecological niche.

Most of our knowledge regarding the molecular details of the underlying regulatory pathways is based on the analysis of the fungal response to single polysaccharides. While this was important to delineate many of the known signaling components, the heterogeneous nature of lignocellulosic substrates demands an understanding of the molecular interplay between the separate regulatory pathways. Our observations of *N. crassa* growth on complex biomass suggested a relation between the cellulase activity and the mannan content of the biomass. We therefore used genetics, biochemical and rheological approaches to find that mannan and cellulose signaling pathways involve common components and are interconnected. Surprisingly, this cross-talk does not lead to synergies but rather leads to confusion on the molecular level with negative effects on cellulase production in several tested fungi. This study thereby provides insights that advance our fundamental understanding of the complex network behind the cross-talk between regulatory systems governing plant cell wall perception and can potentially be applied to produce industrially favorable fungal strains with a lower propensity to be inhibited in presence of complex biomass.

## Results

### The presence of mannodextrins inhibits *N. crassa* growth on cellulose

Comparing the cellulase activity of *N. crassa* WT growing on different carbon sources, we initially observed a consistently lower enzymatic activity on softwood-derived wood powders as carbon source than on hardwood-derived materials and grasses (Fig. 1A). A compositional analysis verified the main difference between hardwoods and softwoods being the content of hemicelluloses. Hardwoods usually have higher xylan content while the main hemicellulose in softwoods are mannans (Fig. S1A) [8,32,33]. We hypothesized that the higher amount of mannan present in softwoods might be involved in the inhibition of cellulase activity of *N. crassa*. To verify this hypothesis, we utilized a biochemical genetics approach aiming to provoke a stronger effect of mannan by artificially altering its intracellular metabolism. The genome of *N. crassa* encodes only one gene (NCU00890) encoding a predicted β-mannosidase for the processing of (gluco-)mannodextrins into monomers [34], a member of the glycosyl hydrolase family two (GH2-1) with no predicted N-terminal secretion signal peptide [35]. To this end, we checked the cellulosic activity of both the WT and the GH2-1 deletion strain (Δ*gh2-1*) grown on the same complex carbon sources as used above. The Δ*gh2-1* strain showed a sharp decrease in total cellulase activity which correlated well with an increased mannan content of the biomass, suggesting a connection between both parameters with a half maximal inhibitory concentration (IC50) of about 1.1% of mannan (Fig. 1B, Fig. S1A-C). To further verify this result, we grew WT and Δ*gh2-1* on mannan-free bacterial cellulose (Fig. S1F) [36] and added low concentrations (0.03% (w/v) corresponding to 3% (w/w) of the used bacterial cellulose) of commercially available mannans or mannobiose to roughly mimic the mannan content present in softwood (Fig. S1A). The added mannan and even mannobiose was sufficient to inhibit cellulase production in the WT and provoked an even more severe phenotype in the Δ*gh2-1* strain (Fig. 1C). To directly test which sugar molecules may cause the inhibition, both the WT and Δ*gh2-1* mutant strain were grown on Avicel, a mannan-contaminated microcrystalline cellulose (Fig. S1F) [36–38]. Afterwards, the HSQC spectra for the anomeric region of the extracted intracellular sugars of both strains were observed by NMR (nuclear magnetic resonance). In comparison to WT, the Δ*gh2-1* strain was found to accumulate mannose as part of a β-1,4-polymer, glucose as part of a β-1,4-polymer, and reducing-end β-mannopyranosyl in the cytosol (Fig. 1D). These results provide strong evidence for β-1,4-linked (gluco-)mannodextrins being the causative molecules for the observed inhibition.

**Fig 1.**
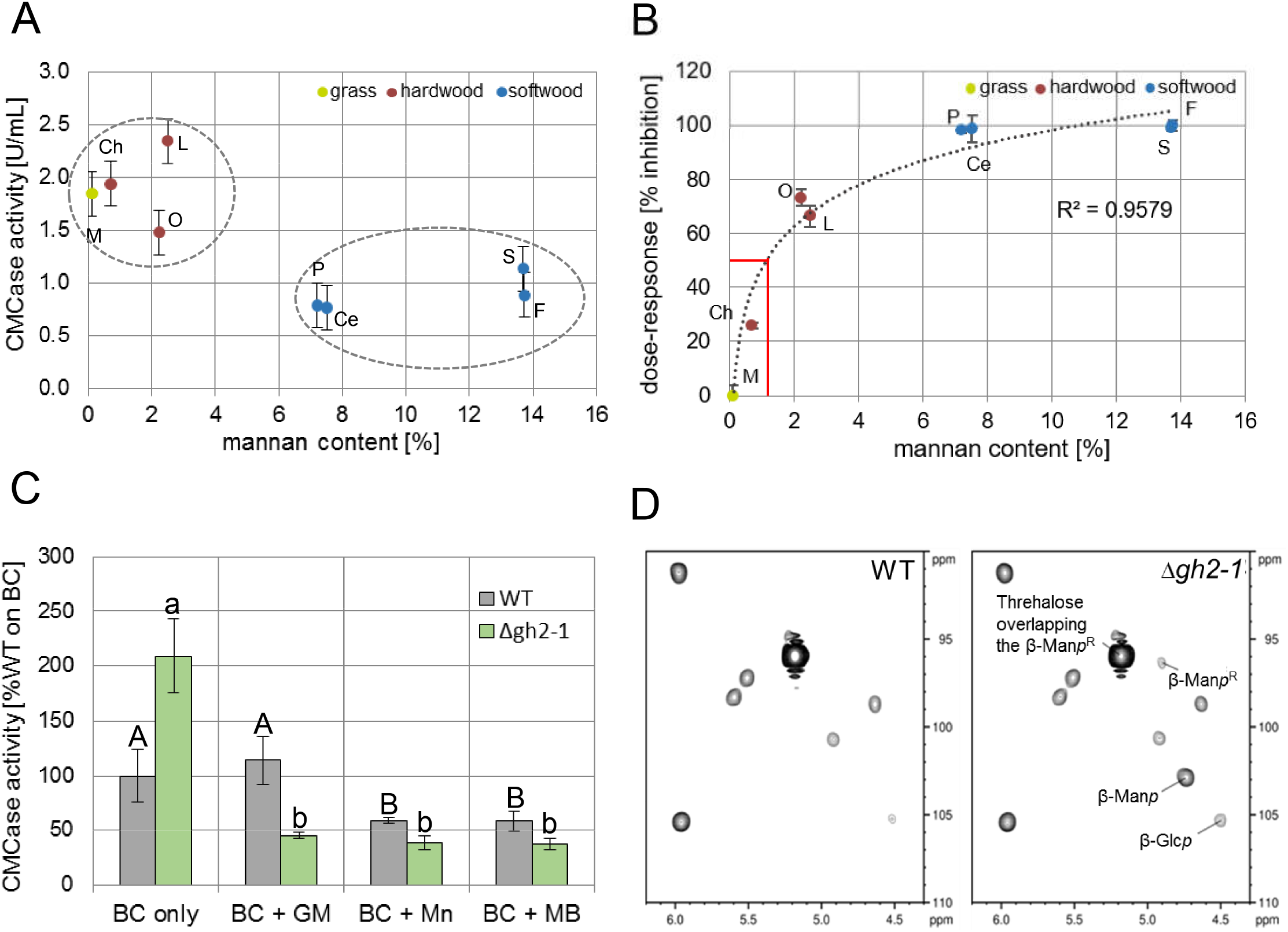
High mannan content is inhibitory for cellulase activity. **(A)** CMCase activity of enzymes secreted into WT culture supernatants after 3 days of growth in 1% (w/v) powdered biomass (M: *Miscanthus*, Ch: chestnut, O: oak, L: locust, P: pine, Ce: cedar, S: spruce and F: fir). **(B)** CMCase activity of the WT and Δ*gh2-1* cultures after growth the same as in (A). **(C)** CMCase activity of the WT and Δ*gh2-1* cultures after growth in 1% (w/v) bacterial cellulose (BC) with the addition of 0.03% (w/v) glucomannan (GM) and mannan (Mn) or mannobiose (MB). **(D)** NMR analysis of the HSQC spectra for the anomeric region of the extracted intracellular sugars of the mycelia of both WT and the Δ*gh2-1* strains after growth in 2% (w/v) Avicel for 24 h post-transfer. β-Glcp: glucose as part of β-1,4-polymer, β-Manp: mannose as part of β-1,4-polymer, and β-ManpR: reducing-end β-mannopyranosyl. Different lower and upper case letters indicate data groups that are significantly different (one-way ANOVA, p-values < 0.05 were considered significant).

### Characterization of the predicted β-mannosidase

To verify its predicted function, GH2-1 was heterologously expressed and purified. The purified enzyme showed strong activity on *ρ*NP-β-D-mannopyranoside with high specificity compared to its activity on *ρ*NP-4-Nitrophenyl-β-D-cellopyranoside, *ρ*NP-4-Nitrophenyl-β-D-glucopyranoside, and *ρ*NP-4-Nitrophenyl-α-D-mannopyranoside as substrates (Fig. 2A). Also a GFP-fusion construct displayed cytosolic localization *in vivo* (Fig. S1D). When assayed at a combination of different temperatures and pHs, in parallel, GH2-1 showed the highest activity in a temperature range between 43 and 54 °C and a pH range between 6.25 and 7.5 (Fig. 2B) and a thermostability up to about 49 °C (Fig. S1E). Moreover, to assess the possibility of mannodextrin cleavage by cross-reactivity of β-glucosidases, we tested the hydrolysis of *ρ*NP-β-mannopyranoside by cytosolic protein extracts from WT, Δ*gh2-1*, Δ*3βG* (a strain carrying deletions for all three β-glucosidase genes [10]) and Δ*qko* (the Δ*3βG* strain crossed to Δ*gh2-1*) grown on 1% Avicel. Only strains possessing GH2-1 displayed β-mannopyranosidase activity (Fig. 2C). Also, when complementing the Δ*gh2-1* strain with the *gh2-1* gene under control of its native promoter and terminator, gh2-1-comp, it showed WT-like β-mannosidase (Fig. 2C). Assaying the cellulase production by the WT, Δ*gh2-1*, and gh2-1-comp strains grown in 1% Avicel showed that in addition cellulase inhibition was relieved in the gh2-1-comp strain (Fig. 2D). This indicated a functional complementation of the *gh2-1* deletion mutation. In summary, these assays confirmed that GH2-1 is the main cytosolic hydrolase encoded in the *N. crassa* genome capable of cleaving mannodextrins.

**Fig 2.**
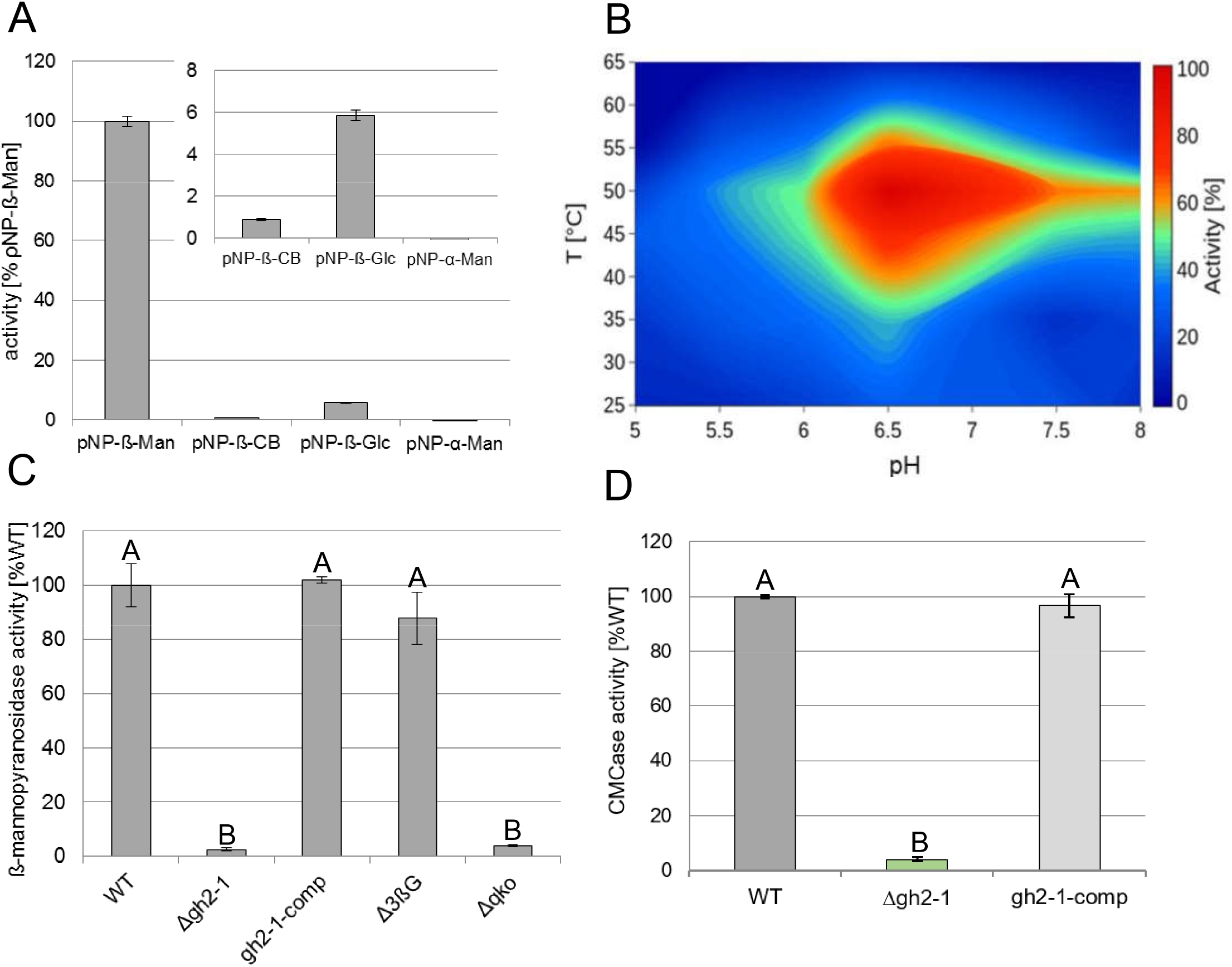
Characterization of GH2-1. **(A)** Substrate specificity assay of GH2-1 using ρNP-4-Nitrophenyl-β-D-mannopyranoside (ρNP-β-Man), ρNP-4-Nitrophenyl-β-D-cellopyranoside (ρNP-β-CB), ρNP-4-Nitrophenyl-β-D-glucopyranoside (ρNP-β-Glc) and ρNP-4-Nitrophenyl-α-D-mannopyranoside (ρNP-α-Man) as substrates. **(B)** Contour plot for GH2-1 activity, at different combinations of temperatures and pHs in parallel, using ρNP-4-Nitrophenyl-β-D-mannopyranoside as a substrate. **(C)** β-mannopyranosidase activity of the cytosolic protein extracts of the WT, Δ*gh2-1*, Δ*3βG* and Δ*qko* (the Δ*3βG* strain crossed to Δ*gh2-1*) after growth in 1 % (w/v) Avicel with 1x Vogel’s salts for 3 days. **(D)** CMCase activity of the WT, Δ*gh2-1*, and *gh2-1-comp* cultures after growth in 1% (w/v) Avicel with 1x Vogel’s salts for 3 days. Different lower and upper case letters indicate data groups that are significantly different (one-way ANOVA, p-values < 0.05 were considered significant).

### A delicate intracellular balance between cello- and mannodextrins

Considering the substantial inhibition caused by intracellular accumulation of mannodextrins, the question arose whether this could be the result of a possible conflict with cellulose signaling. We therefore wanted to assess the influence of the intracellular cellodextrin levels on cellulase inhibition in the Δ*gh2-1* strain. To this end, a cross with Δ*gh1-1*, a deletion strain of the main intracellular β-glucosidase gene [10] was created. Deleting *gh1-1* in the Δ*gh2-1* background completely rescued the Δ*gh2-1* phenotype on mannan-contaminated cellulose (Avicel) (Fig. 3A). This indicated that the effect of accumulating mannodextrins could be counterbalanced by raising the intracellular concentration of cellodextrins. Since the presence of cellodextrins leads to the induction of CLR-2 via the activation of CLR-1 [31], we next tested the possibility to suppress the inhibited phenotype of Δ*gh2-1* by constitutive expression of *clr-2*, rendering its protein levels independent of the levels of its inducing molecules (strain Δ*gh2-1 clr-2 oex*; [31]). Indeed, inducer-independent overexpression of *clr-2* was able to (partially) rescue the Δ*gh2-1* phenotype on Avicel (Fig. 3A).

**Fig 3.**
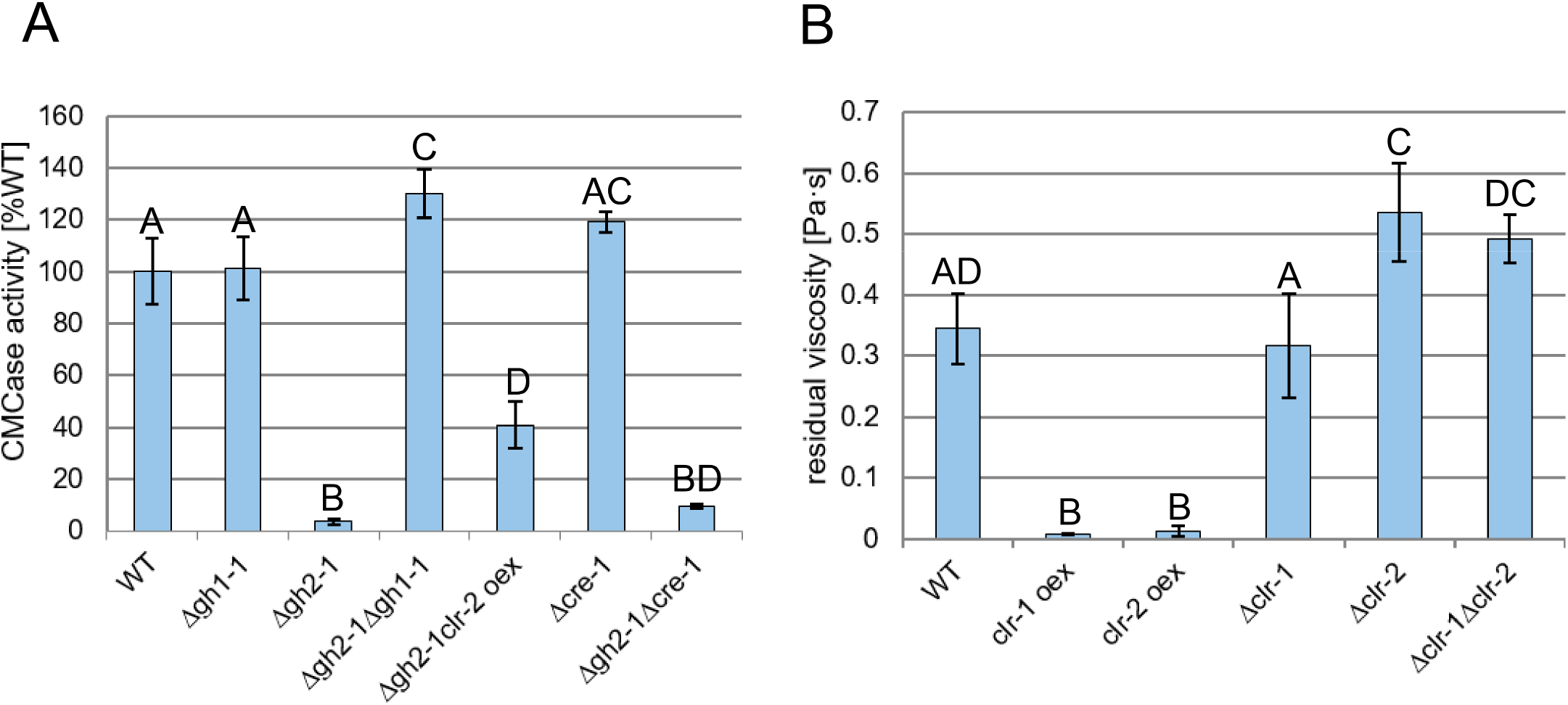
Cello- and mannodextrins compete intracellularly, and the inhibition is independent of CCR by CRE-1. **(A)** CMCase activity of culture supernatants of the indicated strains after growth for 3 days in 1% (w/v) Avicel. **(B)** Viscosity of the culture supernatant of the indicated strains 8 h post-transfer to 1% (w/v) glucomannan. Different lower and upper case letters indicate data groups that are significantly different (one-way ANOVA, p-values < 0.05 were considered significant).

The accumulation of polysaccharide degradation products in Δ*gh2-1* could theoretically also have led to activation of CCR. We thus tested the possibility that the observed inhibition might be due to repression by CRE-1 and studied the effect of a *cre-1* deletion on the phenotype. However, a de-repression due to the loss of CRE-1 in the Δ*gh2-1* background did not lead to a substantial relief of inhibition when grown on 1% Avicel (Fig. 3A), arguing against an involvement of CCR.

Taking into account that CLR-2 is an ortholog of ManR, the regulator of mannan degradation in *A. oryzae* [12,18], and that ChIP-seq data showed CLR-2 to be a direct regulator of *gh5-7* [14], the main predicted β-mannanase-encoding gene in *N. crassa*, we hypothesized that the regulatory pathway of mannan perception shares a common ancestor with the cellulolytic one. This led us to grow *clr-1* and *clr-2* deletion (Δ*clr-1*, Δ*clr-2* and Δ*clr-1* Δ*clr-2*) and mis-expression (*clr-1 oex* and *clr-2 oex*) strains on glucomannan as sole carbon source. By measuring the culture viscosities over time, we aimed to detect the decrease in molecular weight of the hemicellulose polymer [39] due to mannanolytic degradation. Besides *clr-2 oex* also the *clr-1 oex* strain led to a significantly stronger decrease in glucomannan viscosity than the WT strain (Fig. 3B), indicating an enhanced enzyme production on this substrate, which in the case of *clr-1 oex* however might have been an indirect effect via CLR-2. On the other hand, Δ*clr-2* and Δ*clr-1* Δ*clr-2* strains showed a significantly lower reduction in glucomannan viscosity (Fig. 3B), suggesting that CLR-2 is indeed involved in the regulation of mannan degradation in *N. crassa*.

### Cello- and mannodextrins also compete at the level of uptake

Since our data strongly indicate that mannodextrins are cleaved into their constituent monosaccharides only intracellularly by GH2-1, we investigated the transport of mannodextrins into the cell. The two MFS-type transporters CDT-1 and CDT-2 are known to facilitate the uptake of both cellodextrins and xylodextrins [40,41]. Due to structural similarity of (gluco-)mannodextrins, we hypothesized that CDT-1 and −2 might be involved in the uptake of mannodextrins as well. To this end, we tested the growth of the individual and double knockout strains (Δ*cdt-1*, Δ*cdt-2* and Δ*cdt-1* Δ*cdt-2*) in 1% glucomannan. The individual deletion strains for *cdt-1* and *cdt-2* had 66.5% and 85.5% biomass compared to the WT strain, respectively. More significantly, the Δ*cdt-1* Δ*cdt-2* strain had a biomass reduction of about 51% compared to WT (Fig. 4A), indicating an involvement in metabolism of glucomannan. We next tested whether the loss of either CDT-1 or CDT-2 would lead to an impaired uptake of mannobiose by *N. crassa*. For this, sucrose pregrown cultures ofWT, Δ*cdt-1* and Δ*cdt-2* were first induced on 2 mM cellobiose, and then transferred to mannobiose. Following the residual concentration of mannobiose in the culture supernatant, the uptake was found to be almost completely abolished in the Δ*cdt-1* strain (Fig. 4B), whereas its transport was slightly reduced (by about 18 %) in the Δ*cdt-2* strain compared to the WT. We further used *Saccharomyces cerevisiae* that is unable of endogenously transporting cellobiose, to heterologously express CDT-1 or CDT-2[40]. The yeast cells were incubated in either cellobiose or mannobiose for 30 minutes. Indeed, not only cellobiose was imported by both *S. cerevisiae* strains, but also mannobiose (Fig. 4C). Notably, CDT-1 even preferred mannobiose over cellobiose, with only about 18% of mannobiose remaining in the culture supernatant compared to about 40% of cellobiose over the background of empty-vector transformed cells. Moreover, when both sugars were present simultaneously, cellobiose and mannobiose import by CDT-1 was reduced by about 33% and 61%, respectively, indicating that there is a competition between both sugars at the level of uptake by CDT-1 (Fig. 4C).

**Fig 4.**
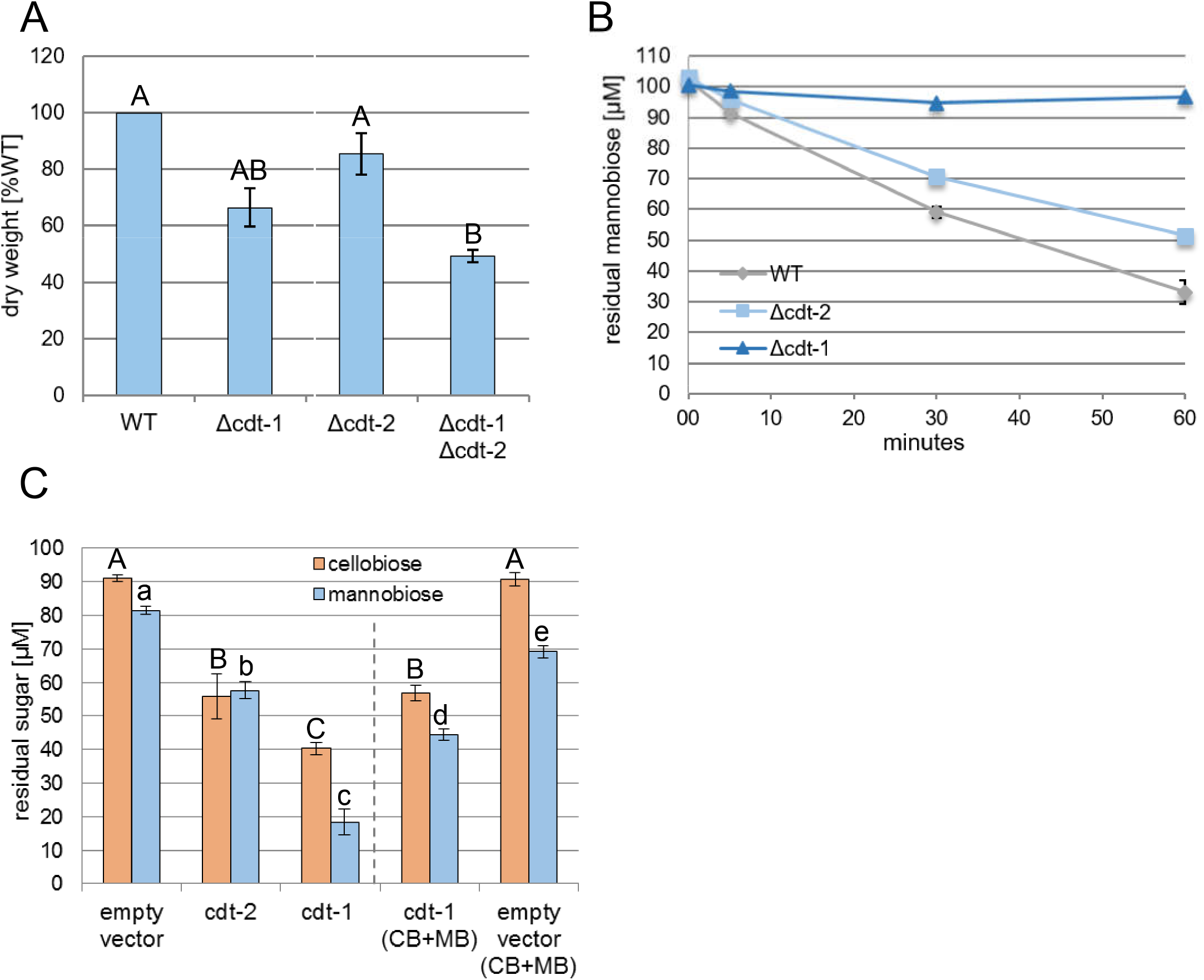
Cello- and mannodextrins compete at the level of sugar uptake. **(A)** Mycelial dry weight of the indicated strains after growth for 3 days in 1% (w/v) glucomannan (indicated in % of WT). **(B)** Residual mannobiose in the supernatant of the indicated strains at indicated times post-transfer to the uptake solution (100 μM mannobiose). **(C)** Residual sugars in the culture supernatants of S. cerevisiae heterologously expressing CDT-1 or −2 transporters, 30 minutes post-transfer to the 100 μM uptake solutions (cellobiose (CB) or mannobiose (MB), or both disaccharides simultaneously). Different lower and upper case letters indicate data groups that are significantly different (one-way ANOVA, p-values < 0.05 were considered significant).

### The inhibitory effect of mannan is conserved in the industrially relevant species *Myceliophthora thermophila* and *Trichoderma reesei*

Lignocellulosic substrates are regularly composed of >1% of mannan. Given the potential impact of the mannan-elicited inhibition on industrial cellulase production, we wanted to test if the inhibition is also present in industrially relevant fungal species. For this, we grew the thermophilic fungus *M. thermophila* [42] on 1% hardwood-derived cellulose being naturally poor in mannan (Emcocel [36]) with and without adding 0.05% glucomannan. Glucomannan addition clearly had an inhibitory effect on cellulase activity (Fig. 5A). Importantly, we checked if the effects are also present for the cellulase hyper-producing *T. reesei* strain RUT-C30.To this end, we grew both *N. crassa* WT and RUT-C30 on 1% Emcocel with and without the addition of 0.05% glucomannan. Similar to *N. crassa*, the low amount of glucomannan was therefore sufficient to significantly reduce total production of cellulases by RUT-C30 (Fig. 5B). This indicates that the overlap between cellulose and mannan signaling pathways appears to be conserved, showing a similar inhibition of cellulase induction in both *M. thermophila* and *T. reesei* as well.

**Fig 5.**
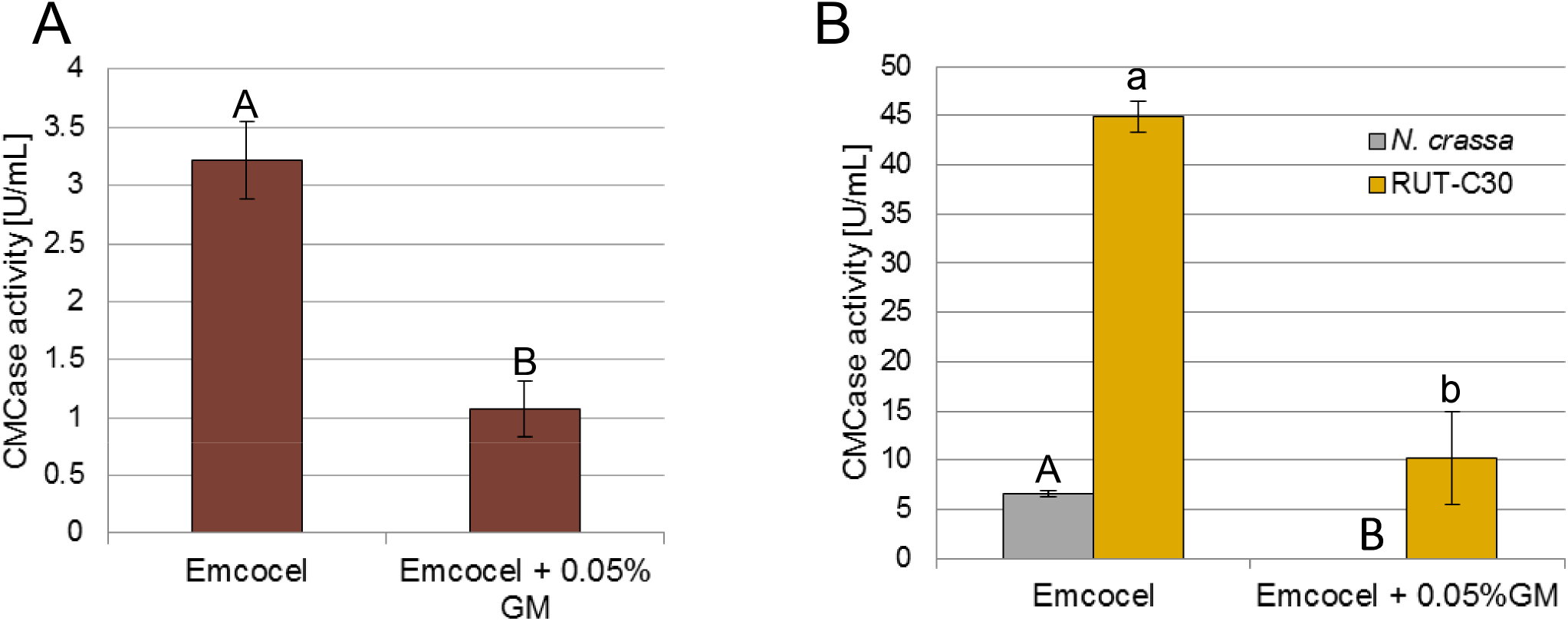
Mannan addition is inhibitory to cellulase production in *T. reesei* and *M. thermophila* as well. CMCase activity of culture supernatants of **(A)** *M. thermophila* WT strain and **(B)** *N. crassa* WT and *T. reesei* RUT-C30 after 3 days growth in 1% (w/v) Emcocel with or without the addition of 0.05% (w/v) glucomannan (GM). Different lower and upper case letters indicate data groups that are significantly different (one-way ANOVA, p-values < 0.05 were considered significant).

## Discussion

In the current model of plant cell wall degradation by *N. crassa*, starvation will lead to the production of low quantities of polysaccharide-degrading enzymes and sugar transporter genes to degrade potential food sources present in the surrounding environment [43]. When cellulose and glucomannan are present (Fig. 6A), secreted cellulases and mannanases degrade the polysaccharides into smaller cellodextrins (such as cellobiose) and mannodextrins (such as mannobiose), which are transported into the cytosol via sugar transporters. While it is known that cellodextrin transporters CDT-1 and CDT-2 transport cellodextrins [40], we found that both transporters are capable of transporting mannobiose as well (Fig. 4). This provides evidence that both transporters are also involved in hemicellulose perception and transport, which is in line with the xylodextrin transport activity already found for CDT-2 by Cai *et al*. [41]. Using *S. cerevisiae* as heterologous expression system, we were able to show that both molecules compete at the level of transport by CDT-1, which even prefers mannobiose over cellobiose (Fig. 4C). Cellobiose and mannobiose have a similar intramolecular β-(1,4)-glycosidic bond between their respective hexose units [44], and their constituent sugars (d-glucose and d-mannose, respectively) are C-2 epimers. Taking this into account, it might be possible that their structural similarity allows them to interact with the same transporters. This is in line with previous findings showing different sugar transporters to have broader specificities by which they have the ability to transport several structurally related sugars. For instance, the *N. crassa* transporters GAT-1 and XAT-1 are capable of transporting galacturonic/glucuronic acid [45] and D-xylose/L-arabinose [46], respectively. Also, the transporter encoded by *mstA* was shown to transport d-xylose, D-mannose and D-glucose in Aspergillus [47], and the fungal D-fructose permease RhtA also accepts L-rhamnose (both 6-deoxy-hexoses) [48].

**Fig 6.**
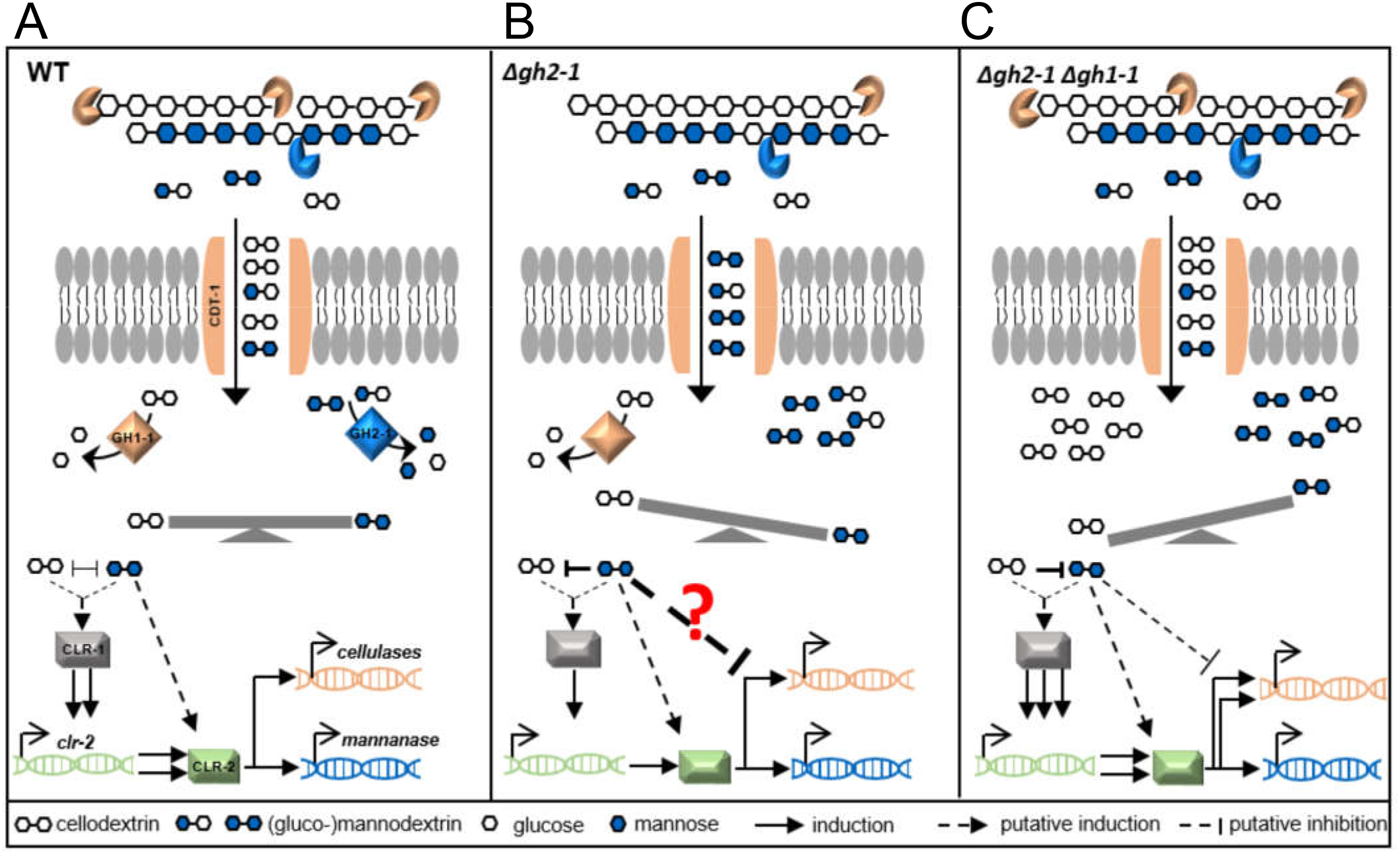
A model of the induction (A), inhibition (B) and relief of inhibition (C) of cellulase production in *N. crassa*. After the degradation of cellulose and glucomannan by cellulases (endo- and exo-acting glucanases, orange) and mannanase (blue), respectively, (gluco)mannodextrins outcompete cellodextrins extracellularly at the level of transport by the (MFS)-type transporter CDT-1. Intracellularly, cellodextrins and (gluco)mannodextrins are further cleaved into the corresponding glucose and mannose monomers by the action of the intracellular β-glucosidase (GH1-1) and β-mannosidase (GH2-1), respectively. In case of an intracellular balance between cello- and mannodextrins (A), an unknown signaling cascade will lead to the activation of the upstream transcription factor CLR-1, which induces expression of the downstream transcription factor CLR-2, which then evokes the major cellulolytic and mannanolytic responses. In the Δ*gh2-1* deletion strain (B), undigested (gluco)mannodextrins accumulate in the cytosol disrupting the intracellular balance of signaling molecules and outcompeting the positively inducing cellodextrins, leading to a potential “confusion” in the cellulolytic signaling pathway in a sense that the fungus is unable to determine the “adequate” amount of cellulase enzymes to be produced, eventually causing a reduced cellulase production. When *gh1-1* is deleted in the Δ*gh2-1* background (Δ*gh2-1* Δ*gh1-1* strain, C), the accumulating mannodextrins could be counterbalanced by the higher amount of undigested cellodextrins present in the cytosol, which re-inforce the induction of the cellulolytic response and relieve the inhibition.

A multitude of previous and ongoing studies have been focusing on understanding the induction of fungal cellulase production by soluble sugars. Many oligosaccharides, such as cellobiose in *N. crassa*, sophorose and lactose in *Trichoderma* and xylose in the Aspergilli, have been identified as inducers of cellulase production [10,49–52]. Yet little is known about an oligosaccharide to have a direct inhibitory effect on the production of such enzymes. *N. crassa* degrades cellodextrins and mannodextrins further into glucose and mannose monomers by the action of (intra- and extracellular) β-glucosidases and the intracellular β-mannosidase GH2-1 (Fig. 6A) [10,11]. Our results indicate that the deletion of this β-mannosidase gene leads to the accumulation of substantial amounts of undigested (gluco-)mannodextrins in the cytosol of *N. crassa*. Our data furthermore provide evidence that these (gluco-)mannodextrins are causative for the strong repression of growth seen for example on mannan-contaminated Avicel as demonstrated by the observation that the addition of mannobiose to *N. crassa* cultures growing on pure bacterial cellulose (as in previous studies [53]) was sufficient to recapitulate the response of the Δ*gh2-1* deletion strain to Avicel (Fig. 1D). Considering the structural similarity between cello- and mannodextrins, their competition at the level of uptake via CDT-1 and the fact that mannodextrins can also inhibit cellobiohydrolase [54], it appears likely that they can also “be confused” by a (yet unknown) signaling component or receptor protein in the cell being somewhat unspecific. The accumulation of (gluco-)mannodextrins is possibly skewing the original balance of signaling molecules in the cytosol and outcompeting the cellodextrins (Fig. 6B). While these would be positively inducing, the interaction with (gluco-)mannodextrins however seems to be unproductive. Likely, this is causing antagonistic effects preventing the native response to cellobiose and interfering with the molecular events leading to the induction of cellulases by the major cellulolytic regulator CLR-2. Generally, less cellulolytic activity results in less substrate degradation and thus lower availability of carbon source and inducing molecules (cellobiose). Eventually, this vicious circle leads to a strong overall signal loss and inhibition of cellulase production and growth (Fig. 6B).

Our use of viscosity measurements as a sensitive tool to detect glucomannan degradation [55] showed that CLR-2 indeed regulates glucomannan degradation, corroborating earlier findings by different methods [14,19]. For instance, ChIP-Seq had identified the genes encoding the β-mannosidase *gh2-1*, the endo-mannanase *gh5-7* and the cellodextrin transporter *cdt-1* to be direct targets of CLR-2 [14]. Homologs of this transcription factor are present in the genomes of many filamentous Ascomycetes including *T. reesei, M. thermophila*, and the Aspergilli [12,17]. In *A. oryzae*, ManR was described to regulate both cellulolytic and mannanolytic genes including the genes coding for the orthologs of the β-mannosidase *gh2-1*, the endo-mannanase *gh5-7* and the cellodextrin transporter *cdt-1* [56]. A similar regulon was also determined for ClrB, the ortholog in *A. nidulans* [31]. These results suggest that the dual function of CLR-2/ManR/ClrB as a combined mannanolytic and cellulolytic TF is conserved from the Aspergilli to *N. crassa*. The role of CLR-2 orthologs in *T. reesei* [20] and *M. thermophila* is much less clear [17]. Nevertheless, the fact that mannodextrins can also induce cellulase inhibition in both strains (Fig. 5) further supports the conserved role of CLR-2. The lack of a clear homolog of CLR-1 in *T. reesei* [20] and the case that an interaction between ClrA and ClrB in Aspergilli may not occur [57], suggest a CLR-1-independent role of CLR-2 and its homologs. This is further supported by the ability of the *clr-1* deletion strain to still utilize glucomannan in contrast to the *clr-2* deletion strain (Fig. 3B). Whether a direct interaction between CLR-2 and both signaling molecules exists or whether these rather interact with an upstream component of the signaling cascade remains to be shown. However, our results support the existence of an intracellular competition upstream of CLR-2, since a misexpression of CLR-2 was able to at least partially rescue the inhibited phenotype (Fig. 3A).

Importantly, we show that there is a delicate intracellular balance between cellobiose and mannobiose which appears to be essential for a full level of cellulase production. While the accumulation of mannodextrins inside the cell has a repressing effect, as seen above, slowing down catabolism of the cellodextrins in the double deletion strain Δ*gh2-1* Δ*gh1-1* counteracts the repression and restores a better cellulosic activity of cellulases (Fig. 6C), presumably by raising the intracellular concentration of cellodextrins. This supports the necessity of a balance that affects the signaling pathway eventually leading to induction or repression of cellulases as presented in our model (Fig. 6).

The ability of molecules to induce or to repress the production of cellulases and hemicellulases by fungi might be masked by CCR. In the Δ*gh2-1* Δ*cre-1* strain, the major TF mediating CCR, *cre-1*, is deleted [30]. Nevertheless, the unaltered inhibition of cellulases caused by the intracellular accumulation of mannodextrins in this strain and the fact that glucomannan was able to inhibit growth on cellulose also in the carbon catabolite de-repressed industrial strain *T. reesei* RUT-C30 [29] confirms that this mannodextrin inhibition pathway is a novel process which is independent of CRE-1-induced CCR during lignocellulose degradation. Moreover, it is known that inducers/repressors are mostly smaller units (oligomers or monomers) that are derived from the polysaccharide itself, such as downstream metabolites and trans-glycosylation derivatives [49,58]. However, in the *N. crassa* Δ*gh2-1* mutant, the β-mannosidase, which would be the likeliest enzyme to perform trans-glycosylation [59], was deleted, suggesting that trans-glycosylation of the inhibitory mannodextrins is not a relevant step for the inhibition.

Although the degradation of cellulose and hemicellulose by filamentous fungi has been intensively studied on well defined, individual polysaccharides, only recently common components for cellulose and mannan perception pathways were described [18,19]. The conservation of the common signaling pathway components (CLR-2, GH2-1 and CDT-1) described in this study suggests that the molecular communication between cellulose and mannan utilization regulatory pathways is likely similarly conserved among filamentous fungi. Moreover, the presence of common signaling intermediates is probably a reflection of the environmental niche of plant-cell-wall degrading fungi, in which cellulose and mannan naturally co-exist in nature [60,61], which allows the fungus to utilize both via common routes.

Finally, taking into account that the industrial production of cellulases is usually performed in presence of residual mannan (either as part of complex plant cell walls or in commercially available plant biomass-derived substrates such as Avicel), this study provides new targets for the improvement of industrial strains for higher cellulase production through the engineering of mannan-insensitivity in the future. This will benefit the development of better enzyme cocktails for the production of biofuels and biochemicals.

## Materials and Methods

### Strains and growth conditions

*N. crassa* strains were obtained from the Fungal Genetics Stock Center (FGSC; [62]) unless indicated otherwise. The Δ*cre-1, 3βG* and *clr-2 oex* strains are a kind gift of N. L. Glass (UC Berkeley, USA). The other knock out strains Δ*qko*, Δ*gh2-1* Δ*gh1-1*, Δ*gh2-1 clr-2 oex*, Δ*gh2-1* Δ*cre-1*, Δ*clr-1* Δ*clr-2* were created through crossings of the respective individual deletion strains as described in the FGSC protocols [62].

All *N. crassa* strains and *T. reesei* RUT-C30 strain (kind gift of M. Schmoll, AIT, Austria) were maintained as described before [36]. *M. thermophila* WT strain (obtained from DSMZ, strain DSM1799) was maintained on 2% (w/v) sucrose Vogel’s minimal medium [63] at 45 °C for 10 days to obtain conidia.

For gh2-1 complementation strain (gh2-1-comp), the *gh2-1* gene amplified from gDNA was placed using SacII restriction site under the control of its native promoter and terminator in plasmid pCSR. The construct was transformed into the Δ*gh2-1* (A) deletion strain by electrotransfection.

The *clr-1* misexpression strain (*clr-1 oex*) was constructed as described in [31] but by using Sbfl and Pacl restriction sites to insert the *clr-1* gDNA in the pTSL126B plasmid placing the *clr-1* gene under the control of the *ccg-1* (clock-controlled gene 1) promoter. A Δ*clr-1* (a) deletion background was used for transformation by electrotransfection. *S. cerevisiae* strain used in this study was D452-2 transformed with pRS316-CDT1, - *CDT2* [64]. Growth was performed as described by [45].

Growth experiments on complex biomasses and on bacterial cellulose were done in 3 mL of 1 × Vogel’s salts plus 1% (w/v) of the corresponding carbon source in 24 deep well plates (at 25 °C, 200 rpm and in constant light). 0.03% (w/v) of commercially available mannans or mannobiose was added to cultures where indicated. Growth experiments for *N. crassa* (at 25 °C and 200 rpm), *M. thermophila* (at 45 °C and 150 rpm), and *T. reesei* (at 30 °C and 200 rpm) were performed in flasks containing 100 mL 1% (w/v) carbon source as described with 1x Vogel’s (*N. crassa* and *M. thermophila*) or 1x Mandels-Andreotti medium [65] in constant light. For inoculation, generally a respective volume of conidial suspension was added after optical density measurements in order to achieve a starting concentration of 10^6^ conidia/mL.

### Biomass and enzymatic assays

For biomass determination, the mycelial mass was dried for 16 h in aluminum pans at 105 °C and measured afterwards. Azo-CMCase activity assays were done according to manufacturer’s protocols (Megazyme, Ireland, S-ACMC), slightly modified according to [36].

For the β-mannopyranosidase activity, *N. crassa* strains were grown in flasks containing 100 mL 1% (w/v) Avicel with 1x Vogel’s (at 25 °C, 200 rpm and in constant light). The mycelia were then harvested by using a Buchner funnel and glass fiber filters, washed 3 times by about 50 mL of 1x Vogel’s solution, then frozen in liquid nitrogen. Frozen mycelia were ground into powder using freezing-milling method. About 250 mg of frozen mycelia were then lysed for protein extraction by adding 750 μl lysis buffer (50 mM Na_3_PO_4_, 1 mM EDTA, 5 % glycerol and 1mM PMSF at pH 7.4). Samples were kept at −20 °C for 30 minutes, and then centrifuged (at 4°C and 1300 rpm for 10 minutes). Protein concentration was measured with Roti^®^-Quant (Carl Roth, K015.1) as described by the manufacturer.

The β-mannopyranosidase activity was then assayed using 4-Nitrophenyl-β-D-mannopyranoside (Megazyme, Ireland, O-PNPBM) as a substrate according to [66] with the following modification: the reaction mixture (containing 50 mM KP buffer at pH 5.5, 80 μg substrate and 1 μg of intracellular protein solution) was incubated at 45 °C for 1 h, then stopped by the addition of 0.5 M Na_2_CO_3_ (pH 11.5). The absorbance was then measured at an OD of 405 nm.

### Compositional analysis

Compositional analysis of biomass was performed as described previously [67].

### Nuclear Magnetic Resonance (NMR) analyses

For the NMR analysis, both WT and Δ*gh2-1* strains were grown in 2% (w/v) sucrose (Sigma Aldrich, S7903) for 16 h then transferred to 2% (w/v) Avicel (PH-101; Sigma Aldrich, 11365) for 24 h. Then the mycelia were collected and their intracellular metabolites were extracted using a protocol modified from Tambellini *et al*. [68]. Briefly, about 500 mg of homogenized mycelia were incubated for 30 minutes on ice with 24 mL cold CH3Cl:MeOH (1:1) and 6 mL dH2O. Samples were centrifuged at 4°C, 4000 rpm for 15 minutes. Supernatants were collected and re-centrifuged for at 4°C, 12000 rpm for 30 minutes. Samples were dried down in a Speed Vacuum concentrator without heating.

Samples were then dissolved in 440 μL D_2_O, 50 μL of Na_2_HPO_4_ and 10 μL DSS (internal standard) yielding clear solutions at approximately 60 mg/mL in 5 mm tubes. 2D NMR spectra were acquired at 25°C on a Bruker AVANCE 600 MHz NMR spectrometer equipped with an inverse gradient 5-mm TXI cryoprobe. Spectra were referenced to DSS at δH 0.00 ppm, yielding HOD resonance at 4.78 ppm. ^13^C–^1^H correlation spectra (HSQC) were measured with a Bruker standard pulse sequence “hsqcetgps¡sp.2”. All the experiments were recorded with the following parameters: spectral width of 16 ppm in F2 (1H) dimension with 2048 data points (TD1) and 240 ppm in F1 (13C) dimension with 256 data points (TD2); scan number (SN) of 128; interscan delay (D1) of 1 sec; acquisition time of 10 h. Assignments of the anomeric signals were assigned based on reference data from the literature. The NMR data processing and analysis were performed using Bruker’s Topspin 3.1 software.

### GH2-1 heterologous expression

For the heterologous expression of GH2-1, the *gh2-1* cDNA was inserted between EcoRI and XbaI restriction sites on the plasmid pGAPZ-B. The construct was transformed into *Pichia pastoris* X-33 strain by electrotransfection according to Invitrogen protocol (https://assets.thermofisher.com/TFS-Assets/LSG/manuals/pgapz_man.pdf). The growth of the transformed Pichia strain and the preparation of cell lysate were done according to the previously mentioned protocol. The cell lysate supernatant was used for GH2-1 purification by immobilized metal-affinity chromatography (IMAC) of the histidine affinity tag [69]. Elution of the enzyme was performed via a pH gradient of 5.5, 5.0, and 4.5 (elution buffer: 50 mM NaH_2_PO_4_, 300 mM NaCl). Protein concentration was measured with Roti^®^-Quant (Carl Roth, K015.1) as described by the manufacturer.

### Substrate specificity of GH2-1

The substrate specificity of GH2-1 was determined by measuring its activities with 4 different substrates: *ρ*NP-4-Nitrophenyl-β-D-mannopyranoside, *ρ*NP-4-Nitrophenyl-β-D-cellopyranoside, *ρ*NP-4-Nitrophenyl-β-D-glucopyranoside, and *ρ*NP-4-Nitrophenyl- α-D-mannopyranoside as substrates (Megazym). The reactions (50 mM KP buffer pH 5.5, 0.1 μg enzyme and 80 μg substrate) were incubated at 37 °C for 1 h, and stopped by the addition of 0.5 M Na_2_CO_3_ (pH 11.5). The absorbance was then measured at an OD of 405 nm.

### Contour plot of GH2-1 activity

The optimal β-mannopyranosidase activity of GH2-1 at different combinations of temperatures and pHs, in parallel, was assayed according to the setup used before [70] with modifications. The reactions consisted of 50 mM KP buffer at different pHs (5, 5.5, 6, 6.5, 7.5, and 8), 80 μg 4-Nitrophenyl-β-D-mannopyranoside (Megazyme, Ireland, O-PNPBM) and 0.0025 μg of purified enzyme. The reactions were incubated for 15 minutes at different temperatures (25, 35, 45, 50, 55, and 65 °C) in a gradient PCR cycler. Then 0.5 M Na_2_CO_3_ (pH 11.5) were added to stop the reaction. The absorbance was then measured at an OD of 405 nm. Blanked measurements were used to generate the contour plot using plotly [71].

### Viscosity measurements

For the Viscosity measurement, the indicated strains were grown in 2% (w/v) sucrose for 16 h then transferred to 1% (w/v) glucomannan. Culture supernatants were collected after 8 h. The viscosity measurements were carried out on an Anton Paar MCR502 rheometer. The control mode feature TruRate^TM^ of the rheometer was enabled during all measurements. Sandblasted parallel plates with a diameter of 25 mm were used and the gap was varied between 0.5 and 1.1 mm, depending on the available amount of the solution. All experiments were carried out at 25 °C. The Peltier hood of the rheometer was used to cover the geometry and the sample. To avoid sample evaporation, the hood was used without applying the internal air circulation and the lower plate was equipped with a solvent trap filled with water, providing an enclosed volume inside the hood. A constant shear rate of 10 sec^−1^ was applied for 100 sec and sampling rate of the measurement was one point/1 sec. The average of the last 10 points was used for the calculation of viscosity.

### Uptake assays

For the yeast-cell based uptake, yeast strain D452-2 cells transformed with pRS316-*CDT1* or -*CDT2* [64] were used. Uptake assays in *S. cerevisiae* and *N. crassa* strains were performed as described before [40,45] with the following modifications: the induction and uptake media contained 1x Vogel’s salts plus 2 mM cellobiose and 0.5x Vogel’s salts plus 100 μM mannobiose, respectively. Samples of the culture supernatants of each strain were taken at the indicated time points (0, 5, 30 and 60 minutes). The samples were centrifuged (at 12000 rpm for 1 minute) and 50 μl of the supernatant was diluted 1:10 with dH2O. Mannobiose concentration was quantified by High Performance Anion Exchange Chromatography coupled to Pulsed Amperometric Detection (HPAEC-PAD) on an ICS-3000 instrument (Thermo Scientific, USA). 25 μL sample was injected onto a Dionex CarboPac PA200 column (3 × 50 mm guard and 3 × 250 mm analytical columns) and eluted at 30 °C using a gradient of 50-170 mM sodium acetate in 0.1 M NaOH at 0.4 ml/min over 12 minutes.

### Statistical analyses

Experiments were done in biological triplicate, and statistical significance was determined by applying analysis of variance followed by a Tukey test using the statistical computing software R [72]. Values of bars and lines in bar and line graphs, respectively, are the mean of the biological replicates, and error bars in all figures are SDs (n=3).

## Acknowledgements

We thank Petra Arnold, Nicole Ganske and Nadine Griesbacher (HFM, TUM) for excellent technical assistance. We furthermore thank Stefan Bauer and Ana B. Ibáñez (EBI, UC Berkeley) for help with the compositional analyses. We are grateful to Monika Schmoll (AIT Austrian Institute of Technology, Vienna, Austria) for the gift of *Trichoderma reesei* strains, to Jamie H. D. Cate (UC Berkeley) for the yeast strains, and to N. Louise Glass (UC Berkeley) for *N. crassa* Δ*cre-1*, Δ*3βG* and clr-2 oex strains. We are also grateful to Dr. Andreas Weiss (J. Rettenmaier and Söhne) for the gift of the Emcocel HD90. We furthermore thank Stefan Bauer and Chris Somerville (EBI, UC Berkeley) for helpful discussions and critical reading of the manuscript.

## Author contributions

L.H., C.T. and J.P.B. designed research; L.H., L.L., H.S., T.G., and J.P.B. performed research; N.G., C.T. and J.P.B. contributed reagents/analytic tools; L.H., L.L., H.S., T.G., N.G., C.T. and J.P.B. analyzed data; and L.H. and J.P.B. wrote the paper. All authors have read and acknowledged the final version of the manuscript.

## Supporting Information

### S1 Methods

#### Thermal stability assay

The thermal stability assay was carried out by incubating the enzyme at different temperatures (0, 25, 37, 45, 55, and 65°C) for 1 h. Afterwards, the residual activity of 0.1 μg enzyme was assayed with 50 mM KP buffer (pH 5.5) and 80 μg 4-Nitrophenyl-β-D-mannopyranoside substrate (Megazyme, Ireland, O-PNPBM). The reaction was incubated at 37 °C for 5 minutes, then stopped by the addition of 0.5 M Na_2_CO_3_ (pH 11.5). The absorbance was then measured at an OD of 405 nm.

#### Microscopy

For gh2-1-gfp strain, the *gh2-1* gene amplified from gDNA was placed under the control of the constitutive promoter *ccg-1* (clock-controlled gene 1) using Xbal and BamHl restriction sites in plasmid pCCG-C-Gly-GFP. The construct was transformed into the WT *his-3^−^* strain by electrotransfection.

GFP fluorescence was visualized using an epifluorescence microscope with a 100x oil-immersion objective.

**Fig S1.**
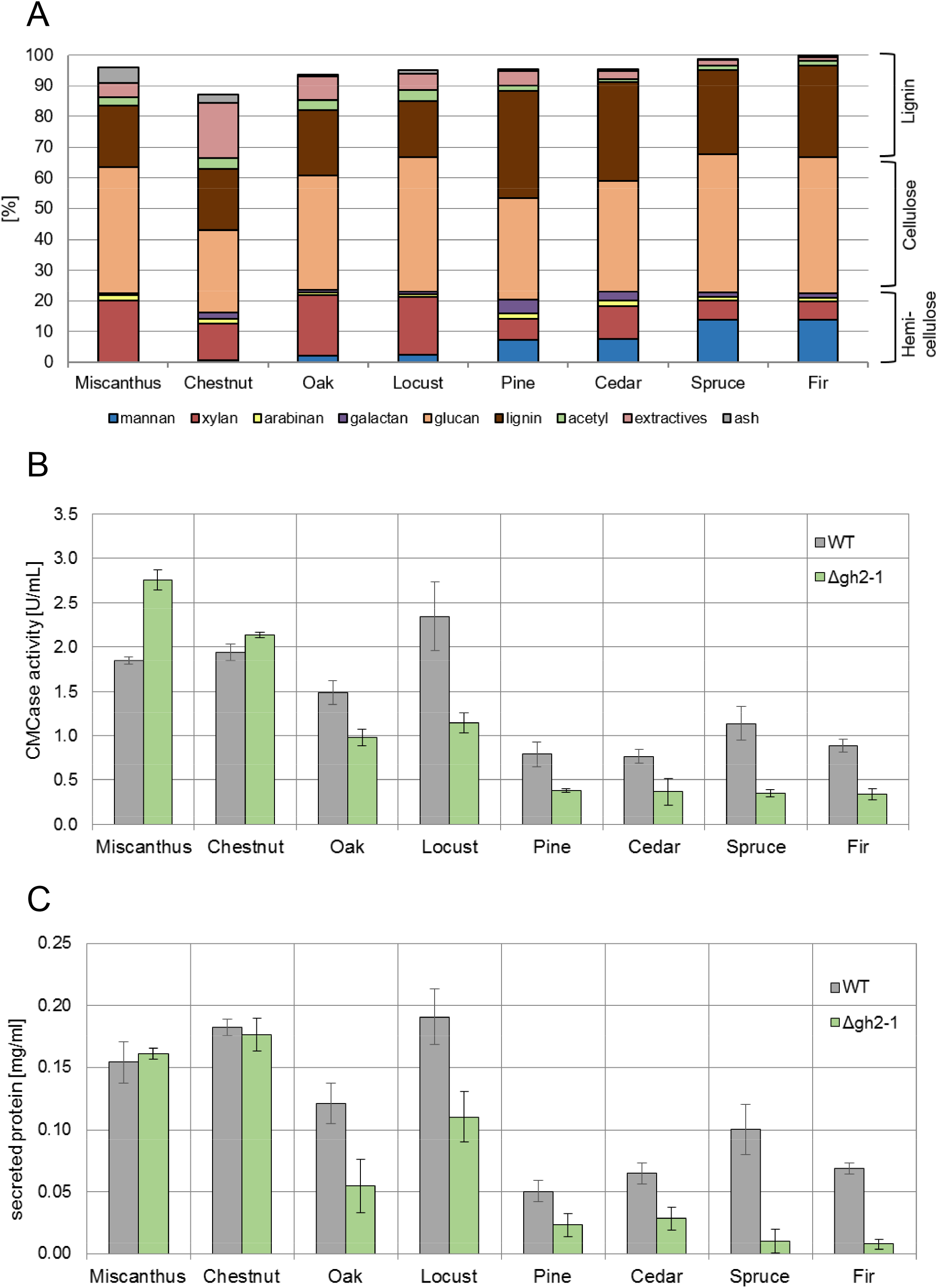
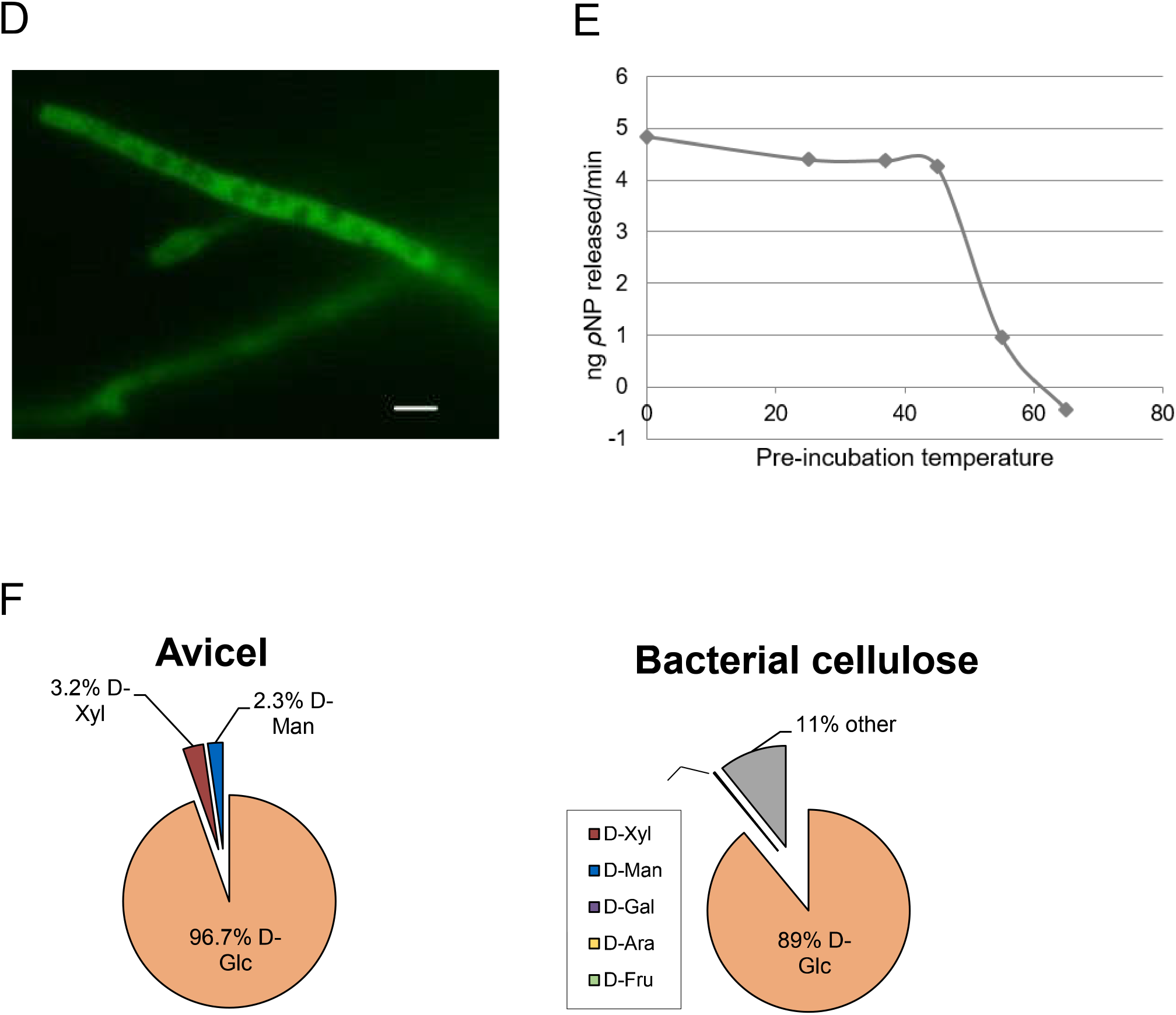
**(A)** Results of compositional analysis of the complex carbon sources (grass: Miscanthus, hardwood-derived: chestnut, oak, and locust, softwood-derived: pine, cedar, spruce and fir) after sulfuric acid hydrolysis (in %). **(B, C)** WT and Δ*gh2-1* phenotype. Both strains were grown for 3 days in 1% (w/v) powdered biomass from different sources with 1x Vogel’s salts (B) then the endo-glucanase activity and (C) the protein concentration of culture supernatants were assayed. **(D)** GH2-1 intracellular localization. A *gh2-1-gfp* strain was used for the localization and visualization by fluorescence microscopy. Scale bar represents 10 μm. **(E)** Thermal stability assay of GH2-1. Purified enzyme was pre-incubated for 1 h at the indicated temperatures, and then the β-mannopyranosidase activity was assayed at 37 °C for 5 minutes. **(F)** Results of compositional analysis of Avicel and bacterial cellulose after sulfuric acid hydrolysis (in %). Glc: glucose, Xyl: xylose, Man: mannose, Gal: galactose, Ara: arabinose, and Fru: fructose.

